# Serotonin buffers hippocampal neuroplasticity and emotional behavior

**DOI:** 10.1101/123646

**Authors:** Giacomo Maddaloni, Sara Migliarini, Francesco Napolitano, Andrea Giorgi, Daniele Biasci, Alessia De Felice, Alberto Galbusera, Sara Franceschi, Francesca Lessi, Marco La Ferla, Paolo Aretini, Chiara Maria Mazzanti, Alessandro Gozzi, Alessandro Usiello, Massimo Pasqualetti

**Affiliations:** Department of Biology, Unit of Cell and Developmental Biology, University of Pisa, 56127, Pisa, Italy.; Ceinge Biotecnologie Avanzate, 80145, Naples, Italy.; Department of Molecular Medicine and Medical Biotechnology, University of Naples “Federico II”, Naples, Italy.; Statistics and Computational Biology Group, Cancer Research UK Cambridge Institute, Li Ka Shing Centre, Cambridge, UK.; Functional Neuroimaging Laboratory, Istituto Italiano di Tecnologia, Center for Neuroscience and Cognitive Systems @ UniTn, 38068, Rovereto, Italy.; Fondazione Pisana per la Scienza, 56100, Pisa, Italy.; Department of Environmental, Biological and Pharmaceutical Sciences and Technologies, Second University of Naples (SUN), Caserta, Italy.; Neuroscience Institute, National Research Council (CNR), 56124, Pisa, Italy.

## Abstract

Adaptation to environmental insults is an evolutionary mechanism essential for survival. The hippocampus participates in controlling adaptive responses to stress and emotional state through the modulation of neuroplasticity events, which are dysregulated in stress-related neuropsychiatric disorders. The neurotransmitter serotonin (5-HT) has been proposed as a pivotal player in hippocampal neuroplasticity in both normal and neuropsychiatric conditions though its role remains still poorly understood. Here, we investigated the impact of 5-HT deficiency on hippocampal activity combining RNA-seq, *in vivo* neuroimaging, neuroanatomical, biochemical and behavioral experiments on 5-HT depleted mice. We unveil that serotonin is required for appropriate activation of neuroplasticity adaptive mechanisms to environmental insults. Bidirectional deregulation of these programs in serotonin depleted mice is associated with opposite behavioral traits that model core symptoms of bipolar disorder. These findings delineate a previously unreported buffering role of 5-HT in instructing hippocampal activity and emotional responses to environmental demands.

---

The hippocampus plays a principal role in stress response and emotional behavior (Kim and Diamond, 2002; McEwen et al., 2015). Changes in hippocampal neuroplasticity are critical in emotional regulation and response to changing environment (Pittenger and Duman, 2008). Growing evidence indicates that adaptive mechanisms to stressful situations involve neuroplasticity events, required to process new information coming from the environment and to organize an appropriate behavioral response (Magarinos et al., 2011; Gray et al., 2014; Gold, 2015). By contrast, disrupted hippocampal neuroplasticity, in terms of reduced neurotrophic support (Autry and Monteggia, 2012), dampened adult neurogenesis (Snyder et al., 2011) dendritic atrophy and spine loss (Watanabe et al., 1992; Pawlak et al., 2005) is thought to underlie maladaptive emotional response to chronic environmental insults recapitulating core symptoms of stress-related neuropsychiatric disorders (Lupien et al., 2009).

The involvement of 5-HT in emotional regulation is well established (Nutt, 2002). The levels of 5-HT, the cellular mechanism for its reuptake/degradation and the activity of serotonergic neurons are reported to be dysregulated in animal models of mood disorders (Bekris et al., 2005; Bambico et al., 2009; Crawford et al., 2010). Importantly, several hippocampal neuroplasticity mechanisms disrupted in these pathological conditions are normalized upon serotonergic drug treatment (Santarelli et al., 2003; Autry and Monteggia, 2012; Boldrini et al., 2012; Morais et al., 2014). However, the direct role of 5-HT signaling in the hippocampus remains unclear. Here, we investigate the impact of 5-HT deficiency on hippocampal transcriptome and its functional activity in *Tph2* -/- mice, depleted of brain 5-HT.

We performed total RNA sequencing to measure the global gene expression in the hippocampus of *Tph2* -/- mice (KO) and we identified 123 differentially expressed (DE) genes in 5-HT depleted mice relative to wild-type (WT) controls, with a p-value adjusted for false discovery rate (p_adj_) ≤ 0.05 (Fig. 1A). DE genes were enriched in the Gene Ontology (GO) categories of ion transport, neurogenesis, neuron projection, and regulation of cell proliferation (Fig. 1B). Interestingly, 5-HT deficiency was associated with increased expression of immediate-early genes (*Arc, Nr4a2, Egr2, Ier5*), ion channels (*Scn4b, Trpv4, Trpm3*), as well as genes involved in neuroplasticity (*BDNF, Nr4a2, Bmp6, Bmp7, Vgf, Enpp2, Igfbp2, Otx2, Calr, Capn6, Folr1, Ttr, Adora2a*, Ace) and neurotransmission (*Nptx2, Slc6a12, Slc6a13;* Fig. 1A). In contrast, the expression of chloride and potassium channels (*Ano2, Kcnn3*) as well as of other genes involved in inhibitory pathways (*Gabra2, Sstr2, Shisa9*) was decreased (Fig. 1A). Such a transcriptional signature led us to hypothesize a tonic hyperactive state in the hippocampus of KO mice.

**Fig. 1:**
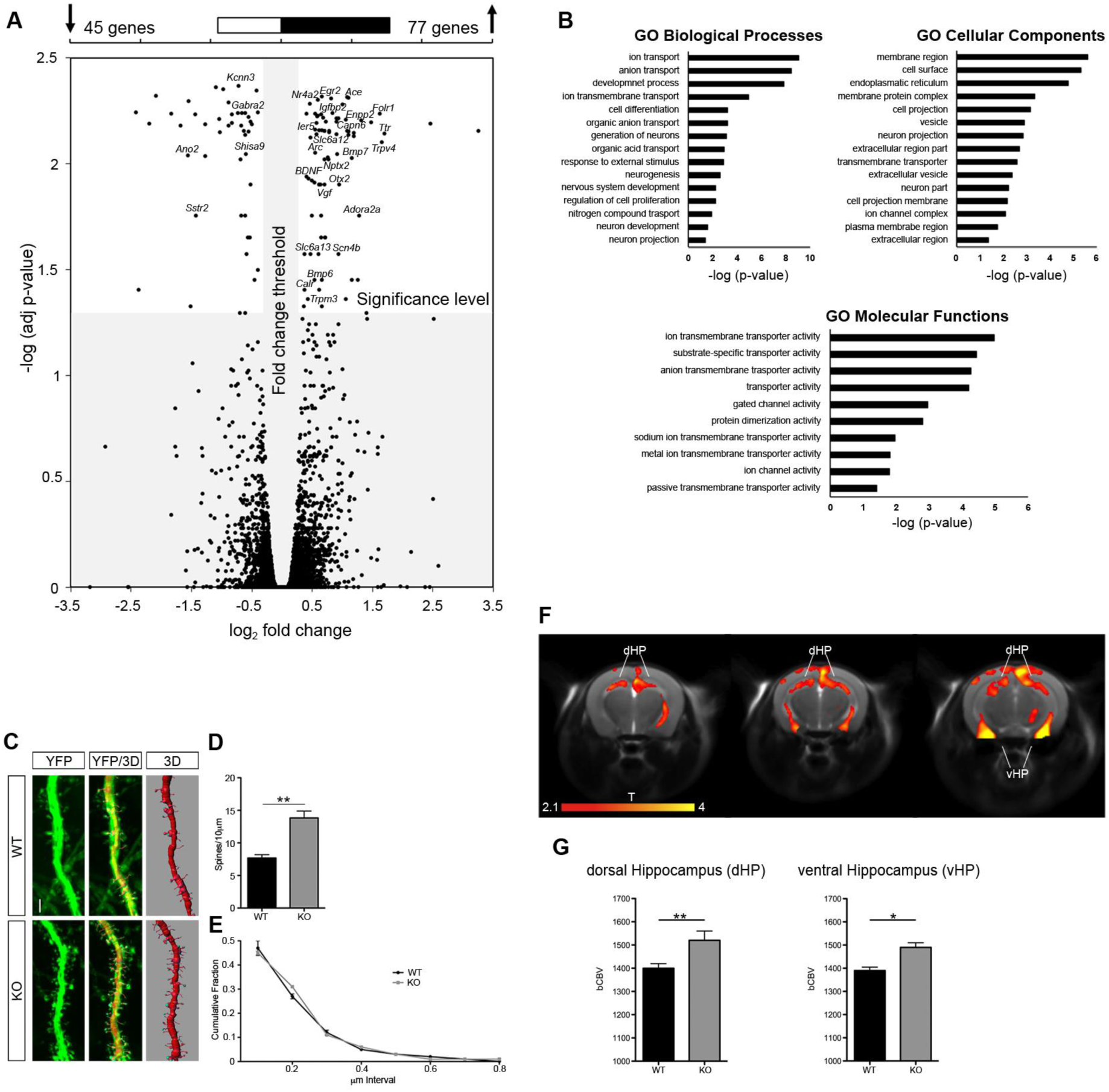
Hippocampal hyperactivity in 5-HT depleted mice. (**A**) Scatterplot of p_a_dj-values versus fold-change between WT and KO mice for all genes. Grey area indicates cut-offs for significance. (**B**) Gene Ontology categories identified from the DE genes. (**C**) Representative images of YFP-labeled CA3 apical dendrites and their 3D reconstruction. (**D**) Quantification of spine density and (**E**) volume distribution in CA3 dendrites (n = 8, for spine density one-way analysis of variance, ANOVA, *F*_*1,13*_ = 18.569). Data are expressed as mean ± s.e.m. **p<0.01. (**F**) Anatomical distribution and (**G**) quantification of brain areas exhibiting a significant increase in bCBV in KO mice with respect to control WT littermates (T>2.1, corrected cluster significance p=0.01). Foci of increased bCBV (red/orange) are superimposed onto contiguous 0.75mm MRI coronal images. The effect has been quantifies in hippocampal areas on slice-to-slice basis. Data are expressed as mean ± s.e.m. *p<0.05, **p<0.01. dHPC, dorsal hippocampus; vHPC, ventral hippocampus. *n* indicates biological replicates. Scale bar = 2μm.

We then examined spine density and morphology along apical dendrites of hippocampal pyramidal neurons in KO adult mice. Spine density along the apical dendritic tree of CA3 pyramidal neurons showed a dramatic increase in KO animals with respect to control littermates (Fig. 1C, D). A trend for a similar effect in CA1 was also apparent (Fig. S1A, B). The distribution of spine volume was otherwise comparable in the two genotypes (Fig. 1E and Fig. S1C). Next, we measured in vivo basal cerebral metabolism by means of basal cerebral blood volume (bCBV) weighted functional Magnetic Resonance Imaging (fMRI) (Gaisler-Salomon et al., 2009; Gozzi et al., 2013), which highlighted clear foci of increased metabolism in hippocampal, but not cortical regions of KO mice (Fig. 1F, G and Fig. S2). Collectively, these data suggest that lack of serotonin leads to aberrant functional hyperactivity in the hippocampus.

We next sought to determine whether this aberrant increased functional activity in the hippocampus of *Tph2* mutant mice might be associated with maladaptive emotional responses. Interestingly, KO mice displayed markedly reduced immobility in both the Forced Swim Test (FST) and the Tail Suspension Test (TST; Fig. 2A, B), indicating reduced depression-like behaviors. Furthermore, mutant mice showed reduced latency to feed in a novel environment in the Novelty-Suppressed Feeding Test (NSF), but comparable weight loss and unaltered 5 min home cage feeding, suggestive of reduced anxiety and increased risk-taking in 5-HT depleted subjects (Fig. 2C). Evidence of robustly increased aggression was observed using a Neutral Arena Aggression Test (NAAT), where KO mice frequently attacked conspecifics with escalated aggression (Fig. 2D). Mutants also showed a more intense exploratory behavior in a Novel Home Cage paradigm (NHC; Fig. 2E) and increased number of rearings relative to controls (Fig. 2F) without showing significant differences in total locomotion (Fig. 2G). Altogether, behavioral testing revealed reduced depression- and anxiety-like behaviors, increased risk-taking, unusual aggression as well as slower locomotor habituation to novelty in KO mice, which together are considered mania-like behavioral phenotypes in rodents (Chung et al., 2014; Logan and McClung, 2016), modeling traits occurring during the manic phase of Bipolar Disorder (BD).

**Fig. 2:**
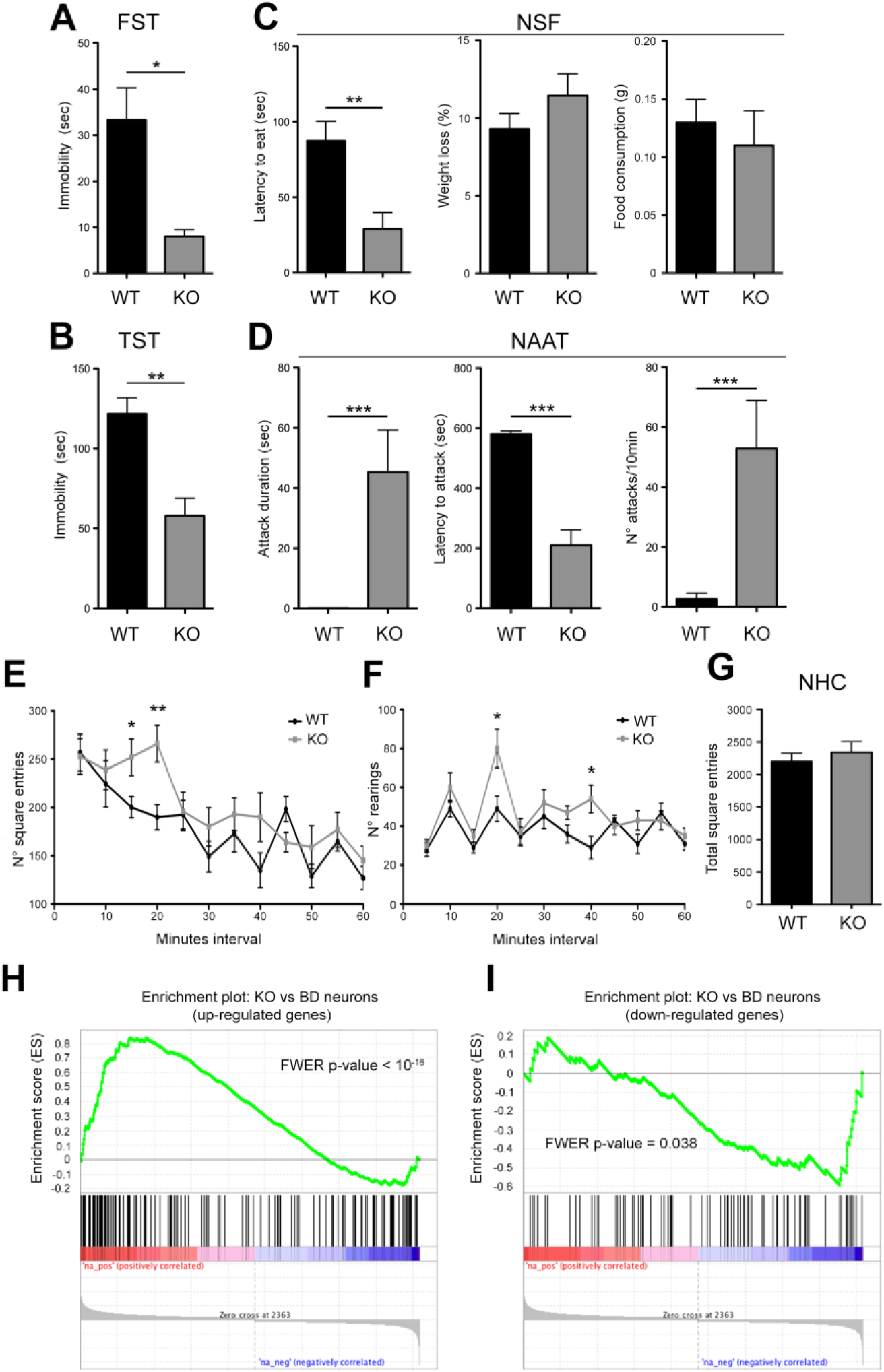
KO mice show mania-like behaviors and common transcriptional abnormalities with neurons derived from BD patients. (**A**) Immobility time in the FST and (**B**) TST. (**C**) Latency to eat, weight loss and food consumption in the NSF. (**D**) Attack duration, latency to the first attack and attack frequency in the NAAT. (**E**) Number of square entries and (**F**) number of rearings in NHC paradigm, respectively. (**G**) Total square entries in the NHC paradigm. (**H** and **I**) GSEA-calculated enrichment between the transcriptional profile of KO mice and that of neurons derived from patients with schizophrenia for (**H**) up-regulated and (**I**) down-regulated genes. For behavioral data, one-way ANOVA followed by Fisher’s post-hoc test were used. FST (*F*_*1,23*_ = 4.317), TST (*F*_*1,17*_ = 8.099), NSF (*F*_*1,23*_ = 8.192), NAAT (duration: *F*_*1,22*_ = 13.073; latency: *F*_*1,22*_ = 49.018; frequency: *F*_*1,22*_ = 15.266), NHC (square entries: *F*_*1,19*_ = 4.519 at 15min; *F*_*1,21*_ = 9.595 at 20min; rearings: *F*_*1,22*_ = 6.182 at 20min; *F*_*1,22*_ = 7.539 at 40min). Data are expressed as mean ± s.e.m., *p<0.05, **p<0.01, ***p<0.001.

The recent evidence that neurons derived from BD patients showed hyper-excitability and increased expression of neurotransmission-related genes (Mertens et al., 2015) led us to investigate potential transcriptional similarities between the gene expression profile of *Tph2* mutants and that of BD neurons. Notably, Gene Set Enrichment Analysis (GSEA) (Subramanian et al., 2005) showed statistically significant enrichment of the KO mouse transcriptional signature as compared to that of BD neurons. Genes up-regulated in KO mice compared to WT were also up-regulated in neurons derived from BD patients compared to healthy controls (Bonferroni corrected p<10^−16^; Fig. 2H), and genes down-regulated in KO mice were also down-regulated in BD neurons (Bonferroni corrected p=0.038; Fig. 2I). Importantly, GSEA did not show any enrichment when the KO mouse transcriptional signature was compared to that of neurons derived from patients affected by schizophrenia (Brennand et al., 2011) (Fig. S3), thus suggesting that the observed enrichment might be BD specific.

BD patients experience abnormal mood transition from the manic to the depressive state. Recurrence of stressful episodes in BD patients overburdens those adaptive mechanisms to stress that are normally recruited in healthy subjects (Kapczinski et al., 2008; Passos et al., 2016), likely promoting pathological mood swings. To assess the possibility to recapitulate such a behavioral transition in 5-HT depleted mice, we exposed WT and KO mice to an 8-weeks unpredictable Chronic Mild Stress (uCMS) protocol and depressive-like behavior was evaluated by means of FST. WT mice subjected to stress (namely WT-S) showed increased immobility relative to WT controls, indicating that uCMS was effective in inducing depressive-like behavior (Fig. 3A). Remarkably, KO mice subjected to stress (namely KO-S) showed a dramatic increase in immobility compared to KO (Fig. 3A, B). We next analyzed hippocampal BDNF/TrkB pathway as it has been reported to be up-regulated 24h after the final stress session in rodents, indicating neuroplasticity events likely involved in adaptive mechanisms to stress (Magarinos et al., 2011; Gray et al., 2014). Results showed in WT-S mice increased BDNF levels 24h after the last uCMS session, whereas KO-S mice displayed reduced expression for both BDNF and TrkB (Fig. 3C, D). These data demonstrate that stress triggers a clear shift to depressive-like phenotype and suggests an impaired mechanism of stress adaptation in *Tph2* KO mice.

**Fig. 3:**
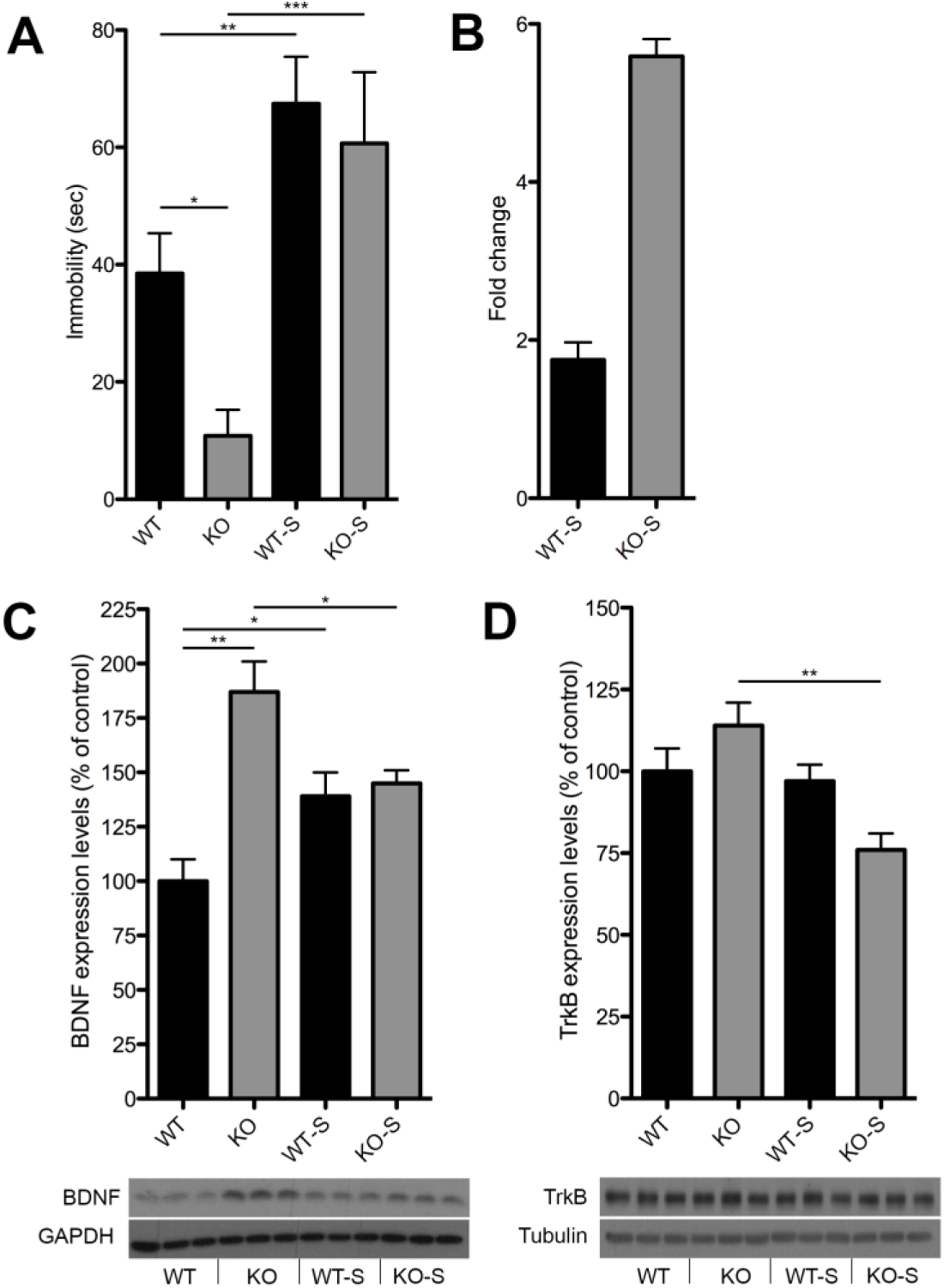
uCMS as a precipitating factor for KO mice behavioral transition. (**A**) Immobility time in the FST of WT-S and KO-S mice and their respective controls (*n* = 20, two-way ANOVA effect of uCMS *F*_*1,72*_ = 14.907 p<0,001; *F*_*1,33*_ = 8.457 WT vs KO: *F*_*1,39*_ = 8,692 WT-S vs WT; *F*_*1,31*_ = 12.825 KO-S vs KO). (**B**) Differential fold-change in immobility time induced by uCMS between the two genotypes. (**C** and **D**) Quantification and representative images of BDNF and TrkB protein levels measured by Western Blotting in WT-S and KO-S mice and their respective controls (BDNF, *n* = 8, two-way ANOVA genotype x uCMS interaction *F*_*1,31*_ = 12.066 p<0,001; Student’s *t* test for single comparisons; TrkB, *n* = 8, two-way ANOVA genotype x uCMS interaction *F*_*1,31*_ = 5.607 p<0.05; Student’s *t* test for KO-S vs KO). Data are expressed as mean ± s.e.m. for behavioral data, as percentage of control ± s.e.m. for biochemical data. *p<0.05, **p<0.01, ***p<0.001. *n* indicates biological replicates.

To explore putative transcriptional changes underlying the mania-to-depression behavioral transition, we performed RNA sequencing on the hippocampus of mice subjected to uCMS. Results showed that stress significantly affected the expression of 144 genes in KO-S mice as compared to KO counterparts (Fig. 4A). Remarkably, 45 of them (e.g. *Otx2, Ttr, Folr1, Enpp2* and *Trpv4*) resulted to be DE in the KO/WT comparison but showing reversed gene expression. This change highlighted a stress-induced lowering of those neuroplasticity factors previously identified in KO mice showing a mania-like phenotype (Fig. 4B, C). Such a unique transcriptional regulation suggested that abnormal variation of neuroplasticity is likely to play a key role in driving opposite behavioral tracts distinctive of BD. Interestingly, further analysis revealed that, out of 140 DE genes identified in the WT-S/WT comparison (Fig. 4A), 53 were DE in the KO/WT comparison (Fig. 4B), all but 1 with the same direction (Fig. 4D). Since these data highlighted an unexpected degree of transcriptional similarity between WT-S and KO mice, we next performed GSEA. Notably, we found a strong significant enrichment between the transcriptional profile of WT-S and KO mice (Bonferroni corrected p<10^−16^ for both up- and down-regulated genes; Fig. 4E, F), suggesting in WT-S mice the establishment of a transcriptional signature that promotes neuroplasticity as a stress-adaptive strategy. These data suggest also that inappropriate activation of the same neuroplasticity factors correlates in the appearance of mania-like behaviors in 5-HT depleted mice. Conversely, 5-HT deficiency blunted adaptive gene programs in the presence of environmental insults (Fig. 4D), demonstrating increased vulnerability to stress in KO-S.

**Fig. 4:**
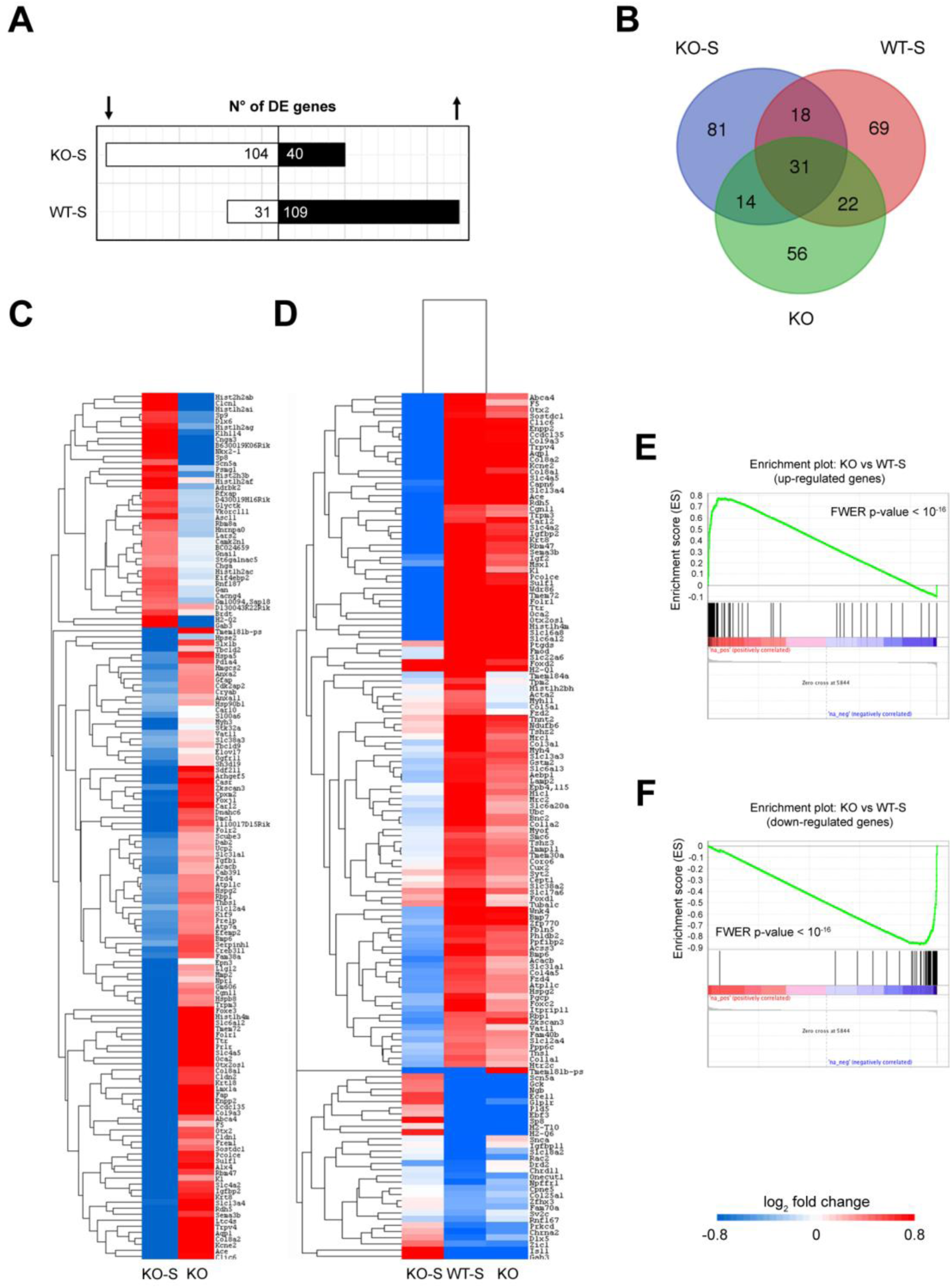
RNA-seq identified an adaptive stress-response gene program modulated by 5-HT that is deregulated in KO mice. (**A**) Schematic representation of DE genes in the WT-S/WT and KO-S/KO comparison. (**B**) Overlapping distribution of DE genes in a Venn diagram across the three comparisons. (**C**) Heat map showing how DE genes in the KO-S/KO comparison (in yellow) are regulated in the KO/WT comparison. (**D**) Heat map showing how DE genes in the WT-S/WT (in yellow) are regulated in the KO-S/KO and KO/WT comparisons. (**E** and **F**) GSEA-calculated enrichment between the transcriptional profile of WT-S and KO mice (ES = 0.77, Bonferroni corrected p<10^−16^ for up-regulated genes; ES = -0.86, Bonferroni corrected p<10^−16^ for down-regulated genes).

Here we demonstrate that 5-HT deficiency results in a transcriptional signature resembling that of human BD neurons and that is associated with increased hippocampal activity, neuroplasticity and mania-like behaviors. Notably, stress reproduced in WT mice a similar transcriptional signature that resulted blunted in KO mice showing a mania-to-depression behavioral switch. On the basis of this evidence we propose that 5-HT signaling in the hippocampus modulates gene programs aimed to promote adaptive responses to stress. Inappropriate activation of these programs might underlie mania-like phenotypes, whereas mania-to-depression transition may arise from the failure to establish an appropriate physiological response.

In conclusion, our findings revealed that 5-HT signaling is central in shaping hippocampal activity and emotional behavior to changing environmental demands, and support the use of *Tph2* -/- mice as a new research tool for mechanistic and therapeutic research in bipolar disorder.

## Materials and Methods

### Animals

*Tph2* -/- mice (Migliarini et al., 2013) and control *Tph2* +/+ male littermates were single-house after weaning in standard Plexiglas cages at constant temperature/humidity (22±1°C, 50-60%) and maintained on a 12/12 h light/dark cycle, with food and water ad libitum. All animals used in each experiment were on a C57BL/6 genetic background. Experimental protocols were conducted in accordance with the Ethic Committee of the University of Pisa and approved by the Veterinary Department of the Italian Ministry of Health.

### RNA extraction

Total RNA was extracted using the automated Maxwell 16 LEV RNA FFPE Purification Kit with the Maxwell 16 Instrument (Promega, Madison, WI, USA). We followed the manufacturer's instructions protocol starting from the Lysis Buffer and Proteinase K step excluding the Mineral Oil procedure. Hippocampus tissue was homogenized in Lysis Buffer using a pestle.

### Whole Transcriptome RNA analysis

RNA-seq was performed using NextSeq 500 (Illumina, San Diego, CA, US) for Next Generation Sequencing. The library was prepared following the protocol TruSeq Stranded mRNA LT kit (Illumina). Libraries were quantified using Qubit 2.0 Fluorometer (Invitrogen, Life Technologies, Grand Island, NY) and the size profile was analyzed on the 2200 TapeStation instrument (Agilent Technologies, Santa Clara, CA).

### RNA-seq data analysis

Raw data were converted to FASTQ format using bcl2fastq (Illumina). We used the FastQC quality control tool (http://www.bioinformatics.babraham.ac.uk/projects/fastqc/) to perform quality assessment. In addition, we evaluated raw data contamination from different organisms (bacteria, fungi, virus) by applying FastqScreen (http://www.bioinformatics.babraham.ac.uk/projects/fastq_screen/). RNA-Seq reads were aligned to the mouse genome (mm10; UCSC) with STAR aligner 2.5.1 (https://github.com/alexdobin/STAR). Differential expression between conditions was calculated using Cuffdiff (http://cole-trapnell-lab.github.io/cufflinks/cuffdiff/). All RNA-Seq analyses were performed in the cloud using the Seven Bridges Genomics platform (www.sbgenomics.com). Gene Set Enrichment Analysis was performed using GSEA (http://software.broadinstitute.org/gsea/). Hierarchical gene clustering on differentially expressed genes was performed using Bioconductor ctc package on R 3.0.1 (https://www.bioconductor.org/packages/release/bioc/html/ctc.html).

### Immunohistochemistry

Animals were perfused transcardially with 4% paraformaldehyde (PFA), brains were dissected, post-fixed o/n at 4°C and sections (50μm thick) were obtained with a vibratome (Leica Microsystems). Immunohistochemistry was performed following standard protocols. Briefly, free-floating sections were permeabilized with 0.5% Triton-X100 (Sigma) in PBS. Sections were then blocked in 5% horse serum (Gibco, Life Technologies), 0.5% Triton-X100 in PBS for 1 h followed by overnight incubation with the primary antibody (rabbit anti-GFP antibody, 1:2000, Molecular Probes) at 4°C. After six washes with 0.5% Triton-X100 in PBS, sections were incubated overnight with the secondary antibody (Rhodamine Red-X goat anti-rabbit IgG, 1:500, Molecular Probes) at 4°C. After three washes with 0.5% Triton-X100 in PBS, section were incubated with DAPI (0.1 μg/ml, Sigma), washed three times with PBS and then mounted onto glass slides and coverslipped with Aqua Poly/Mount (PolyScience).

### 3D modeling analysis

For dendritic spine analysis, adult Tph2 -/- mice and control mice carrying the Thy1-YFP-M allele were processed for immunohistochemistry using anti-GFP antibody and a rhodamine-conjugated secondary antibody to avoid interference of endogenous YFP fluorescence. A number of 6 confocal fields (35 Z-steps at 0.15μ interval) on consecutive coronal sections of both CA1 and CA3 fields in the dorsal hippocampus were imaged using a Nikon A1 confocal microscope equipped with a 60x PlanApo oil objective at 1024×1024 pixel resolution. Images were analyzed using the Filament Tracer semi-automated method for dendrites and spine properties quantifications (Imaris 7.2.3, Bitplane). For each animal, 20 to 30 dendrites in the apical region were reconstructed for each hippocampal field.

### *In vivo* functional Magnetic Resonance Imaging (fMRI)

Animal preparation. Magnetic Resonance Imaging experiments were performed on adult Tph2 +/+ (n = 10) and Tph2 -/- (n = 10) littermate male mice. Briefly, mice were anaesthetized with isoflurane (5%), intubated and artificially ventilated. The left femoral artery was cannulated for contrast agent administration, continuous blood pressure monitoring and blood sampling. At the end of surgery, isoflurane was discontinued and substituted with halothane. Experiments were carried out at a maintenance anesthesia level of 0.8%. Arterial blood gases (paCO2 and paO2) were measured at the end of the functional time series. The values recorded were 16±4 mmHg (paCO2), 287±95 mmHg (paO2) and 17±4 mmHg (paCO2), 272±86 mmHg (paO2) for Tph2 -/- and control, respectively. No significant inter-group difference in paCO2 or (paO2) levels was observed between groups (p>0.65, Student’s t test). Functional data acquisition commenced 30 min after isoflurane cessation.

Image Data Acquisition. All experiments were performed using a 7.0 Tesla MRI scanner (Bruker Biospin, Milan). Transmission and reception were achieved using a 72 mm birdcage transmit coil and a custom-built saddle-shaped solenoid coil for signal reception. Shimming was performed on a 6 mm × 6 mm × 6 mm region, using a FASTMAP protocol. For each session, high-resolution anatomical images were acquired with a fast spin echo sequence (RARE) with the following parameters: repetition time (TR)/echo time (TE) 3550/40 ms, matrix 192×192, field of view 2×2 cm2, 28 coronal slices, slice thickness 0.50 mm. Co-centered cerebral blood volume (CBV) weighted fMRI times series were acquired using a Fast Low-Angle Shot (FLASH) MRI sequence with the following imaging parameters: FLASH TReff = 283.023 ms, TE = 3.1 ms, α=30°; FOV 2 × 2 cm2, 156 × 156 × 500 μm resolution, dt = 60 s, Nr = 60, corresponding to 60 min total acquisition time. Images were sensitized to reflect alterations in CBV by injecting 5 μl/g of superparamagnetic iron oxide (Molday Ion, Biopal) intra-arterially after 5 baseline images.

basal CBV mapping. To calculate basal CBV (bCBV), CBV-weighted time series were spatially normalized to a study-based anatomical template, and signal intensity was converted into basal cerebral blood volume (bCBV(t)) pixel-wise. bCBV time-series were calculated over a 5 minute time-window starting 15 min after contrast agent injection. Voxel-wise group statistics was carried out using FSL using multi-level Bayesian inference and a T threshold > 2.1, and corrected cluster significance threshold of p=0.01.

### Behavioral Testing

All behavioral procedures were performed during the light phase of the cycle (11:00 - 13:00 h). Depression-like behaviors were assessed in the Forced Swim Test (FST) and Tail Suspension Test (TST) that were performed following standard protocols. Briefly, in the FST mice were placed in a 5L Plexiglas Beaker containing 4L of 26°C water and video-recorded for 6 min. Minutes from 2 to 6 were analyzed for immobility time. In the TST, mice were hanged by their tail from a bar 50 cm from the ground with a piece of autoclave tape and were recorded in a 6 min session. Minutes from 2 to 6 were analyzed for immobility time. For both tests, immobility was considered as absence of any active movement of the paws. Anxiety-like behavior was analyzed in the Novelty-Suppressed Feeding (NSF). Food was removed from the cages of mice 24 h before testing. The next day, mice were placed for 10 min in a bright white arena (38×35×20) without bedding with a food pellet at the center. Mice were video-recorded and latency to feed was assessed offline. After the test, to avoid confounding effects of feeding behavior on anxiety, hunger was measured by weighting before and after 5 min a single pellet of food placed in the home cage. Aggressive behaviors were measured in a Neutral Arena Aggression Test (NAAT). Two mice of the same genotype were placed in a novel standard cage (42.5×26.5×18.5cm) with bedding and video-recorded for 10 min. Lateral threats and clinch attacks were considered as sign of aggression. Latency to the first attack, attack duration and number of attacks in 10 min were measured. Locomotor habituation to novelty was measured in the novel home-cage paradigm. Briefly, mice were individually placed into a novel standard cage (42.5×26.5×18.5cm) with bedding and video-recorded for 60 min. The area of the cage was virtually subdivided in squares and the times the testing mouse crossed one of the grid lines with all four paws (i.e. square entry) was scored, as well as the times it stood on its hind legs (i.e. vertical activity, also known as rearing).

### unpredictable Chronic Mild Stress (uCMS)

uCMS paradigm was modified from Tye *et al*. (Tye et al., 2013). Age-matched Tph2 -/- and control mice were randomly subdivided in the uCMS and control group. Control mice of both genotypes were housed in standard conditions. uCMS protocol consisted in two stressors per day (one during the day, one during the night) for 8 weeks. Cage tilt on a 45° angle for 16h, food deprivation for 6h, white noise (http://www.simplynoise.com) for 16h, continuous illumination for 36h, 3h darkness during the light cycle, continuous darkness for 36h, water deprivation for 6h, wet bedding (150mL water into sawdust bedding) for 16h, rat feces exposure in the cage for 16h, cage switching between mice, restraint stress in 50mL tube for 2h, overcrowded housing for 3h were the stressors randomly applied to be unpredictable for mice. Except for overcrowding, as well as for water and food deprivation sessions, water and food were available ad libitum. Tph2 -/- and WT mice from uCMS and control group were behaviorally tested with the Forced Swim Test 24h after the last uCMS session, and then immediately sacrificed. Brains were rapidly removed, hippocampal tissue dissected and quickly frozen in liquid nitrogen for Western blot and RNA-seq analyses.

### Western Blotting

Biochemical studies were performed as reported in Napolitano *et al*. (Napolitano et al., 2010). After dissecting out, the hippocampus were sonicated in a lysis buffer (320mM sucrose, 50mM Tris HCl pH 7.5, 50mM NaCl, 1% Triton X-100, 5mM β-glycerol phosphate, 1mM Na3VO4, 5mM NaF, protease inhibitor cocktail) and incubated on ice for 30min. Samples were spin at 12,000g x 10min and the supernatant transferred to fresh microfuge tube. Aliquots of the homogenate were used for protein determination using Bio-Rad Protein Assay kit (Bio-Rad, Hercules, CA). Equal amounts of total proteins (30 μg) for each sample were loaded onto 15% (for BDNF detection) or 10% (for TrkB detection) polyacrylamide gels. Proteins were separated by SDS-PAGE and transferred overnight to membranes (PVDF; Amersham Pharmacia Biotech, Uppsala, Sweden). The membranes were immunoblotted overnight using selective antibodies against BDNF and TrkB (each diluted 1:1000, Santa Cruz Biotechnology). Both BDNF and TrkB optical density values were normalized using antibodies against GAPDH (1:1000, Santa Cruz Biotechnology) and □-Tubulin (1:50000, Sigma Aldrich), respectively. Blots were then incubated in horseradish peroxidase-conjugated secondary antibodies and target proteins visualized by ECL detection (Pierce, Rockford, IL), followed by quantification by Quantity One software (Biorad). Normalized values were averaged and used for statistical analysis performed by two-way ANOVA followed by post-hoc comparison, when required.

## Acknowledgments

This work was supported by Italian Ministry of Education, University and Research (MIUR) (Prin 2008, 200894SYW2), Toscana Life Sciences Foundation (Orphan_0108 program) and Norvegian Research Council to M.P. G.M. and A. Giorgi were supported by PhD program from University of Pisa. S.M. was supported by Regional Program and European Social Fund. We thank C. Valente for excellent technical assistance and members of our laboratory for valuable discussions and comments on the manuscript.

## Author contributions

S.F., F.L., M.L.F., P.A. and C.M.M. conducted RNA-seq. G.M., A. Giorgi and D.B. performed analysis of RNA-seq data. G.M. and S.M. performed dendritic spine analysis. A.D.F., A. Galbusera and A. Gozzi conducted fMRI analysis. G.M., S.M., F.N., A. Giorgi and A.U. performed behavioral studies. F.N. and A.U. performed Western blot analysis. G.M., S.M., A.U. and M.P. designed the research. G.M., S.M. and M.P. analyzed data. G.M. and M.P. wrote the manuscript.

